# Non-invasive optical synthetization of hearing sensation in non-genetically modified animal

**DOI:** 10.1101/2023.05.13.540641

**Authors:** Yuta Tamai, Miku Uenaka, Aya Okamoto, Keito Hosokawa, Yuki Ito, Koji Toda, Shizuko Hiryu, Kohta I. Kobayasi

**Affiliations:** Neuroethology and Bioengineering, Graduate School of Life and Medical Sciences, Doshisha University, Kyotanabe, Kyoto, Japan; Neurobiology of Social Communication, Department of Otolaryngology-Head and Neck Surgery, Hearing Research Centre, University of Tübingen, Medical Center, Elfriede-Aulhorn-Strasse 5, 72076 Tübingen, Germany; Larning and behavior, Department of Psychology, Keio University, Tokyo, Japan

**Keywords:** auditory perception, brain–machine interfaces, classical conditioning, cochlear implants

## Abstract

Brain-machine interfaces are emerging devices that act as a medium for communication between the neural system and external software and hardware. Over the last decade, there has been a discussion on applying infrared laser stimulation to the brain-machine interfaces, including cochlear implants. Infrared lasers can selectively activate neural populations without introducing exogenous agents to tissues. This study presents the first demonstration that laser irradiation of the cochlea, a peripheral sensory organ, can elicit a clear behavioral auditory response. Mongolian gerbils (*Meriones unguiculatus*) were divided into two groups, each subjected to classical conditioning where a reward water delivery was predicted by either cochlear laser stimulation or sound stimulus. Laser-conditioned animals successfully learned licking behavior, with conditioned responses and behavioral properties comparable to auditory-conditioned animals. The laser-evoked response was significantly inhibited by auditory masking, and auditory-conditioned animals demonstrated stimulus generalization to laser stimulation. In a subsequent experiment, simultaneous presentation of auditory and laser stimulation induced nonlinear amplification in both auditory cortical and behavioral responses, suggesting that combining auditory and laser stimulation can enhance auditory perception beyond the effects of each stimulus alone. These findings indicate that infrared laser irradiation of the cochlea can potentially evoke and enhance auditory perception, making it a promising candidate for implementation in auditory prostheses.

## Introduction

Approximately 430 million people, constituting 5% of the world’s total population, suffer from hearing loss, leading to significant communication challenges and social isolation [1, 2]. For those severely hearing impaired, implantable auditory devices, such as hybrid cochlear implants (which combine hearing aid and cochlear implant technology), represent a limited option [3, 4]. However, these devices require surgical intervention, and 30%–55% of hearing-impaired patients encounter over 30 dB HL of residual hearing loss due to the complications of post-surgical implantation [5]; implantable auditory devices pose challenges, especially in preserving residual hearing for hybrid implants.

The last decade has seen many attempts to apply optical neural stimulation to brain-machine interfaces. The method has the potential to activate spatially selective cell populations in a non-contact manner, and several studies have demonstrated successful optical neuromodulation without invasive probe implantation [6–8]. However, with respect to cochlear implants, most studies have focused on intracochlear optical stimulation to improve the spectral resolution using infrared laser stimulation [9–11] or optogenetic stimulation [12, 13]. Therefore, as far as we know, none has investigated the potential of infrared laser stimulation for minimizing surgical invasiveness, utilizing its non-contact nature.

Here, as the essential foundation for the non-invasive cochlear laser stimulation, we assessed, for the first time, the auditory perceptual event elicited or enhanced by transtympanic laser stimulation. Our prior research utilized the penetrative nature of laser stimulation through thin tissues and demonstrated that transtympanic laser stimulation could elicit cochlear responses [14] and auditory cortical activities [15] bypassing the middle ear. This research systematically characterized laser-induced perception by electrophysiological and behavioral means using Mongolian gerbils, a standard animal model for auditory physiology [e.g., 16, 17, 18]. They were trained to associate a reward (a drop of water, US: unconditioned stimulus) with either auditory, laser, or visual cues (CS: conditioned stimulus) using a classical conditioning paradigm (Fig. 1A–C, Fig. S1 and S2; see Materials and Methods for details). Cochlear responses triggered by laser and auditory stimuli during the conditioning task were recorded to analyze the correlation with behavioral responses. After subject animals were trained with auditory and laser stimuli, we examined the auditory masking effect on the laser-evoked behavioral response and the laser intensity dependence of the behavioral response, and tested stimulus generalization from auditory to laser stimulation. Subsequently, we performed simultaneous presentation of auditory and laser stimulation to evaluate the interaction and the potential for enhancing auditory perception with simultaneous stimulation.

**Fig. 1.**
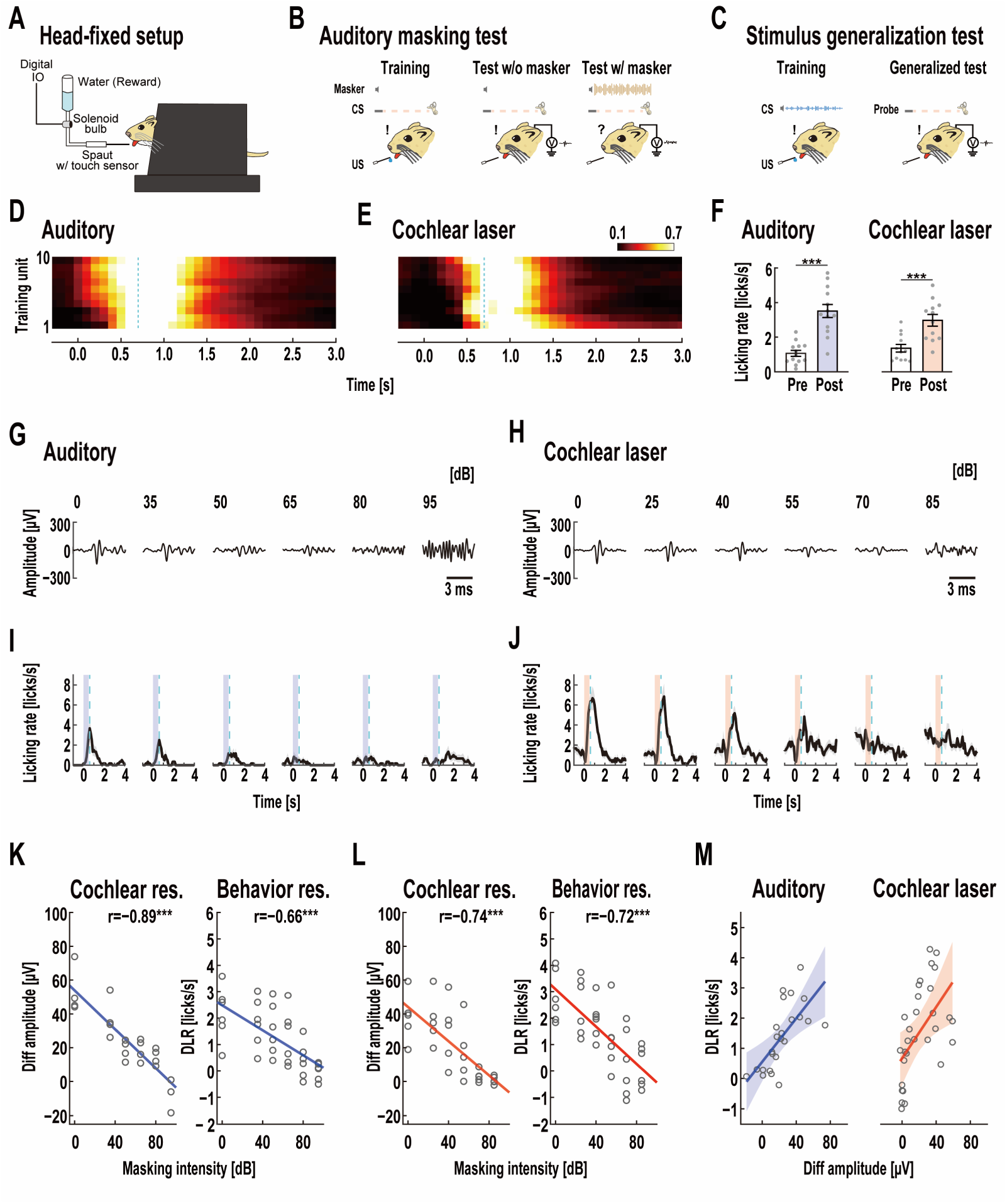
Licking behavior transaction following the training and auditory masking effect on cochlear and behavioral responses evoked by auditory and laser stimulation. (A) Schematic demonstration of the head-fixed setup. Experimental workflow of the auditory masking (B) and stimulus generalization (C) are shown (Materials and Methods for details). Heat map of the auditory-(D) and laser-induced (E) average PSTH of the licking behavior in the rewarded trial of each training unit (auditory: n=13; laser: n=12). Each row is a different training unit. One training unit comprised 100 trials. The blue dashed lines indicate the reward timing. PSTH was standardized at the peak licking rate in each training unit. As the training progressed, the standardized auditory- and laser-induced licking rates were observed earlier by a degree after the CS onset. (F) Mean licking rate with auditory (n=13) and laser (n=12) stimulation on the last training day of catch trials (1–120 trials). The white bars show the mean licking rate for 1 s before stimulus onset. The blue and pink bars indicate the mean licking rate between 0.2 and 1.2 s. The mean licking rates after stimulus onset were significantly higher than those before it in both auditory and laser stimulation. Effect of auditory masking (0, 35, 50, 65, 80, and 95 dB SPL) on CAP from auditory nerves evoked by 80 dB SPL auditory (G) and 13.2 mJ/cm^2^ laser (H) stimulation. Mean PSTH of licking behavior induced by 80-dB SPL auditory (I) and 13.2-mJ/cm^2^ laser (J) stimulation. The blue and red zones in (I) and (J) indicate auditory and laser stimulation periods, respectively. Gray areas show the SEMs of each trial. Light blue dotted lines represent the expected reward time. Changes in different amplitudes of CAP and DLR induced by auditory (K) and laser (L) stimulation with different masking intensities. As the intensity of the masker on auditory-evoked cochlear and behavioral responses increased, CAP amplitude and DLR significantly decreased (CAP amplitude: r=−0.89, \*\*\**P*<0.001; DLR: r=−0.66, \*\*\**P*<0.001; Pearson’s correlation analysis). A similar tendency was observed in laser-evoked cochlear response and behavioral response, where the intensifying sound pressure level of the masker caused a significant decrease in CAP amplitude (r=−0.74, \*\*\**P*<0.001; Pearson’s correlation analysis) and DLR (r=−0.72, \*\*\**P*<0.001; Pearson’s correlation analysis). (M) Linear regression of DLR changes on the diff amplitudes of CAP. Blue and red lines show straight line fits, and blue and red areas show 95% confidence intervals. As an increase in DLR was observed depending on the different amplitudes of CAP in auditory and laser stimulation conditions (auditory: r=0.80, \*\*\**P*<0.001; laser: r=0.81, \*\*\**P*<0.001; Pearson’s correlation analysis). ANCOVA showed no significant difference in slopes (F(1,50)=0.177, *P*=0.676; ANCOVA) and y-axis intercepts (F(1,51)=0.741, *P*=0.393; ANCOVA) of the auditory- and laser-induced relationship between different amplitudes of peripheral neural response and DLR. (F and K to M) Open gray circles show individual data. Further behavioral alteration by Pavlovian conditioning is shown in supplemental figure (Fig. S3).

## Results and discussion

### Transtympanic laser stimulation elicits auditory perception

Subject animals were successfully trained with cochlear laser stimulation as a conditioned stimulus (Fig. 1D–F, and Fig. S3). After approximately 1000 training trials (10 units), all auditory- and laser-trained subjects (auditory: n=13; laser: n=12) could associate the relationship between CS and US. The learning curves of each stimulus were comparable (Fig. 1DE). Additionally, white noise systematically masked both laser-evoked and auditory-evoked behavioral responses but not visually evoked behavioral response (Fig. 1B, G–M and Fig. S4). Previous studies revealed that laser-evoked electrophysiological responses can be masked by acoustic stimulation, indicating that laser-evoked responses are processed in peripheral auditory pathways [19, 20]. Our research has directly evaluated the influence of auditory masking on the laser-induced behavioral consequences. The previous study by Matic et al. [10] investigated laser-induced perceptual events in cats and demonstrated that laser irradiation of the cochlea increased the probability of the animal turning its head to the irradiated side. However, as this response was spontaneous and not associated with specific perception, the content of laser-evoked perception remains unidentified; therefore, our research provides first compelling evidence of auditory perception.

### Auditory perception can be manipulated by changing radiant energy

The amplitude of the laser-evoked compound action potential (CAP) was well-correlated with the strength of the conditioned response (Fig. 2). Although some studies have attempted to show the possibility of manipulating perception in terms of intensity dependency [9, 21] and spectral selectivity [22, 23] by comparing auditory- and laser-evoked physiological responses, no studies have ever revealed the relationship between laser-evoked physiological responses and behavioral consequences. Our data showed that both laser-evoked CAP and behavioral responses increased depending on the radiant energy, which were comparable to those on the sound pressure level (Fig. 2A–F). The same was true in the masking experiment (Fig. 1K-M). These suggest that the laser-evoked behavioral responses were mediated by cochlear activation, and laser-evoked perception can be controlled by changing radiant energy, possibly in a similar way to the sound pressure level of auditory stimulation.

**Fig. 2.**
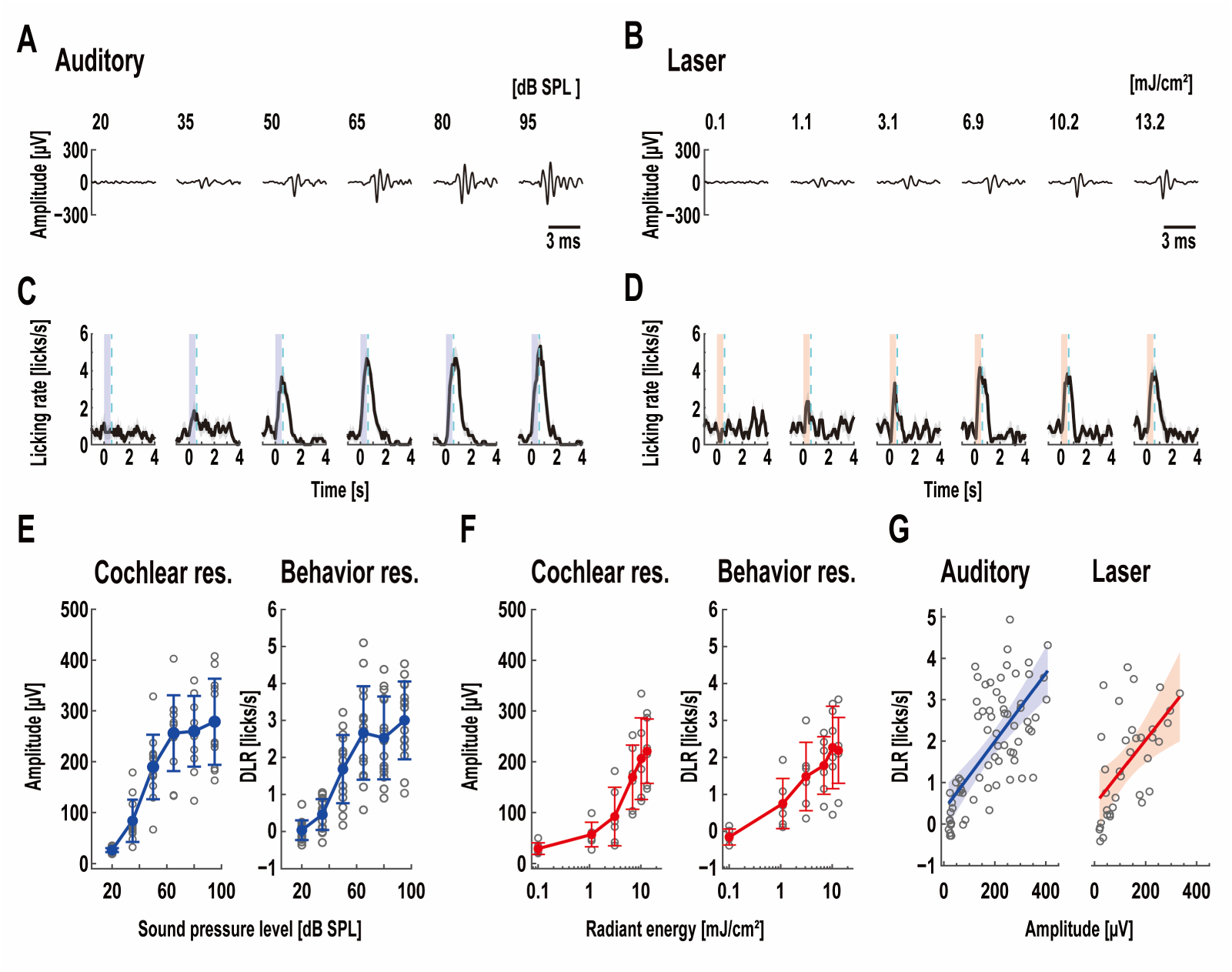
Intensity dependance of cochlear and behavioral responses elicited by auditory and laser stimulation. Compound action potentials (CAP) of auditory nerves with auditory (A) and laser (B) stimulation. Mean peristimulus time histogram (PSTH) of licking behavior elicited by auditory (C) and laser (D) stimulation. In (C) and (D), the blue and red zones indicate auditory and laser stimulation periods, respectively. Gray areas show the standard error of the mean (SEM) for each trial. Light blue dotted lines represent the reward timing in a training session. Increasing the intensity of auditory and laser stimuli produced a systematic change in CAPs and mean PSTHs. Average amplitude of CAP and delta licking rate (DLR) with auditory (E) and laser (F) stimulation. Error bars indicate standard deviations (SDs). As the sound pressure level increased from 20 to 95 dB SPL, the average CAP amplitude and DLR increased from 26.14±3.87 to 278.63±84.83 (n=12; Mean±SD) and 0.03±0.27 to 3.00±1.05 (n=15), respectively. With increasing radiant energy from 0.1 to 13.2 mJ/cm^2^, there was a steady increase in CAP amplitude and DLR (CAP amplitude: from 28.92±11.34 to 220.71±63.38, n=6; DLR: from −0.15±0.21 to 2.19±0.89, n=6). (G) Correlation between CAP amplitude and DLR elicited by auditory and laser stimulation. Blue and red lines depict straight line fits, and blue and red areas show 95% confidence intervals. Increasing CAP amplitude resulted in a systematic increase in DLR under auditory and laser stimulation (auditory: r=0.80, \*\*\**P*<0.001; laser: r=0.81, \*\*\**P*<0.001; Pearson’s correlation analysis). The slopes and y-axis intercepts of the fitted linear regression were not statistically different from each other (slope: F(1, 104)=0.046, *P*=0.832; intercepts: F(1,105)=0.093, *P*=0.761; ANCOVA). (E to G) Open gray circles show individual data. Further intensity dependance of behavioral responses is described in supplemental figure (Fig. S5).

### Laser-induced perception is similar to auditory perception elicited by clicking sound, but at least partially different

To further test laser-evoked perception, we evaluated stimulus generalization in auditory stimulation to laser stimulation (Fig. 1C and Fig. 3). Auditory-conditioned animals consistently exhibited CR after probe laser stimulation, even though the laser was never presented during training (Fig. 3A–H). This result suggests that transtympanic laser stimulation, similar to auditory stimulation, formed acoustic perception. Our data also indicate that increasing the radiant energy of laser stimulation enhances the laser-evoked behavioral response, suggesting that laser irradiation of the cochlea (generalized stimulus) can manipulate auditory perception.

**Fig. 3.**
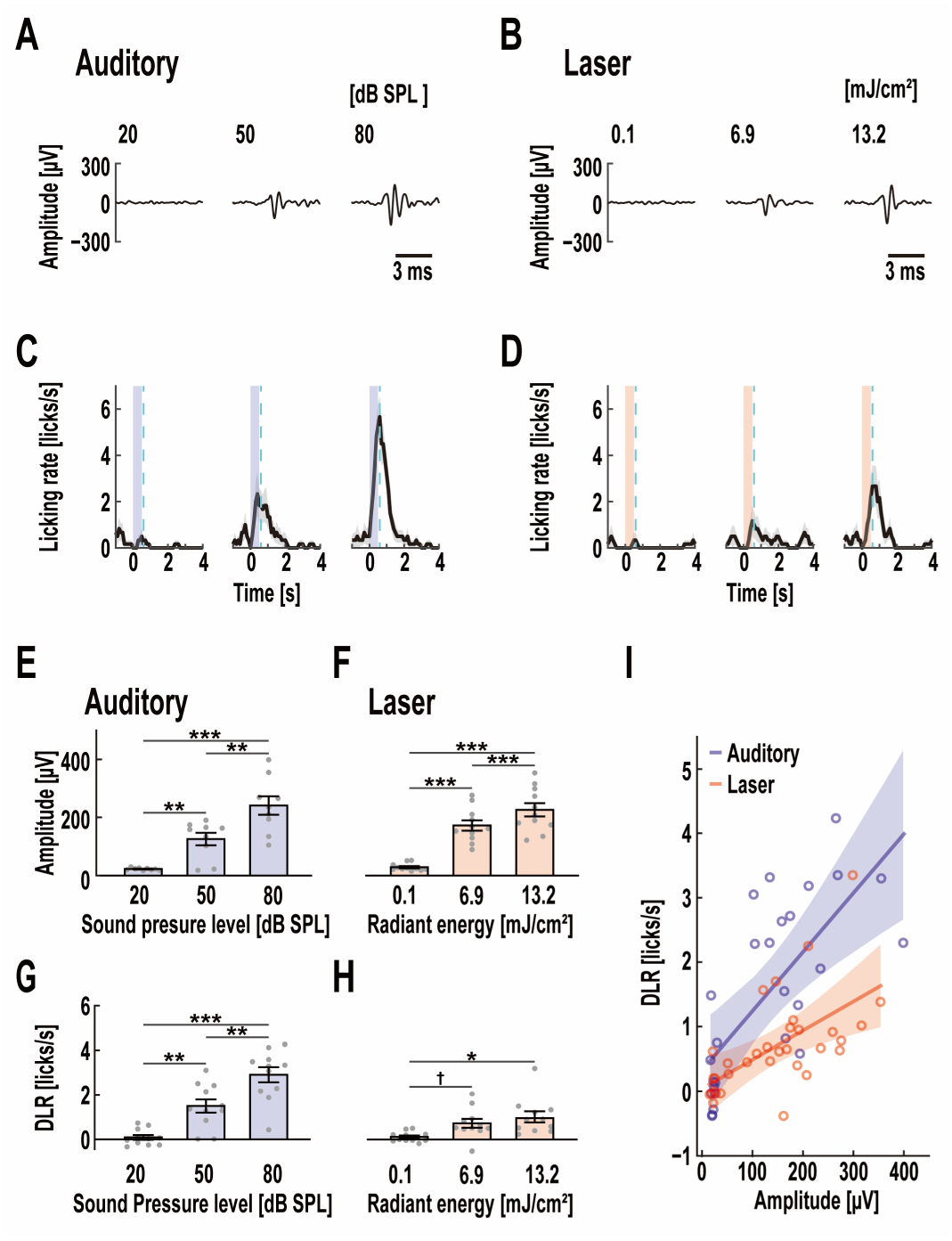
Stimulus generalization from auditory stimulation to laser stimulation. Compound action potential (CAP) from auditory nerves (A and B) and mean PSTH of licking behavior (C and D) elicited by auditory (20, 50, and 80 dB SPL) and laser (0.1, 6.9, and 13.2 mJ/cm^2^) stimulation in subjects classically conditioned with auditory stimulus as CS. The blue and red zones in (C) and (D) indicate auditory and laser stimulation periods, respectively. Gray areas show the SEMs of each trial. Light blue dotted lines represent the reward timing subjects expected. Mean CAP amplitude (E and F) and DLR (G and H) with auditory and laser stimulation. Error bars represent SEMs. Open gray circles show individual data. Auditory- and laser-induced CAP amplitudes significantly increased with stimulus intensity (auditory, 20 dB SPL: 22.45±1.37 µV, 50 dB SPL: 125.33±21.56 µV, 80 dB SPL: 240.79±31.71 µV, n=9; laser, 0.1 mJ/cm^2^: 28.74±3.79 µV, 6.9 mJ/cm^2^: 172.04±17.61 µV, 13.2 mJ/cm^2^: 225.83±22.90 µV, n=11; Mean±SEM). Laser stimulation induced licking behaviors in auditory-trained animals in an intensity-dependent manner similar to auditory stimulation. As the sound pressure level increased from 20 to 80 dB SPL, DLR increased from 0.09±0.11 to 2.90±0.34 licks/s (n=11). Laser-induced DLR gradually changed depending on radiant energy (0.1 mJ/cm^2^: 0.12±0.07 licks/s, 6.9 mJ/cm^2^: 0.73±0.20 licks/s, 13.2 mJ/cm^2^: 1.01±0.25 licks/s, n=11). In (E to H), there was a significant difference in response to each stimulus intensity based on two-tailed paired *t*-test with Bonferroni correction for multiple comparisons (†P<0.1, *P<0.05, \*\**P*<0.01, \*\*\**P*<0.001). (I) DLR as a function of CAP amplitude induced by auditory and laser stimulation. Blue and red lines and shaded areas describe fitted linear regressions and 95% confidence intervals. Open blue and red circles show individual data. The correlation coefficients between CAP amplitude and DLR were statistically significant in both auditory and laser stimulation conditions (auditory: r=0.80, \*\*\**P*<0.001; laser: r=0.81, \*\*\**P*<0.001; Pearson’s correlation analysis). Comparing the linear regression model between auditory and laser stimulation conditions revealed that the difference in slopes and y-axis intercepts was statistically significant (slopes: F(1,56)=5.474, \**P*<0.05; y-axis intercepts: F(1,57)=18.256, \*\*\**P*<0.001; ANCOVA).

However, the detailed features of auditory perception elicited by transtympanic laser stimulation remain unclear. Analysis of covariance (ANCOVA) revealed that the laser-generated linear regression slope in the dependence of DLR on CAP in acoustically trained subjects was significantly smaller than the auditory-generated linear regression slope (Fig. 3I). Conversely, there was no difference between auditory- and laser-generated dependence in subjects trained on each stimulus modality (Fig. 2G). In other words, the laser-induced cochlear response elicited a weaker CR than the auditory-induced cochlear response. However, the amplitude of the laser-induced cochlear response has enough potential to generate a CR comparable to that of the auditory-induced cochlear response. This result suggests that pulsed laser irradiation elicited an auditory perception, albeit at least partially different from auditory-induced perception as a training stimulus (i.e., click train), based on the general insight that CR amplitude peaks at the CS and gradually decreases as the similarity between the CS and unassociated stimuli decreases [24, 25]. Future studies using laser-trained subjects to transfer behavioral responses to various types of auditory stimuli (e.g., different frequency sounds) are required to assess laser-evoked perceptual content in more detail.

### The effect of optoacoustic phenomena in laser-induced auditory response

The mechanism by which cochlear laser stimulation evokes neural and behavioral responses is still largely unresolved. Following the initial study by Wells et al. [26], which demonstrated that laser irradiation of sciatic nerves yielded CAP in vivo, subsequent studies delved into the underlying mechanism of the laser-evoked neural response. These studies suggest that the optical absorption of water causes an insatiable temperature rise in cells [27–29], resulting in reversible alterations in electric capacitance of the plasma membrane [30] or changes in heat-induced ion channel activity [31]. The thermal build-up in cochlear cells also results in basilar membrane vibration with lymph expansion, causing hair cell depolarization (optoacoustic effect) [32–35]. Xia et al. [34] reported that laser stimulation at 13.9–182.3 mJ/cm^2^ might induce basilar membrane vibration corresponding to sound between 46 and 74 dB SPL, and the sound level could be 53.5 dB SPL when laser pulses at 18.9 mJ/cm^2^ (threshold radiant energy) were irradiated to the cochlea. Although our results are not directly comparable to these data due to differences in laser parameters (e.g., spot size, laser irradiation time, and repetition rate), which are challenging to accurately determine in an in vivo experiment, as mentioned by Tan et al. [36], and tissue parameters (e.g., absorption coefficients, density, and thermal conductivity), these results suggest that our laser stimulation potentially induced nonnegligible lymph vibration in the cochlea. Additionally, some studies have shown direct activation of electrophysiological responses in hair cells [37–39]; therefore, laser stimulation has the sufficient possibility of inducing hair cell depolarization.

To explore hair cell function in laser-induced cochlear activation, several studies [35, 40, 41] compared laser-evoked auditory responses in normal hearing and chemically deteriorated cochlea. They demonstrated that laser-evoked cochlear responses were absent in chemically impaired cochlea, suggesting mediation by hair cells. In contrast, Richter and colleagues [42] reported observing the cochlear response induced by laser stimulation after a 30–40 dB chemical deafening of the auditory threshold. They noted that the laser-induced response couldn’t be masked by white noise in animals with chemically deteriorated cochlea [20]. Tan et al. [43] showed that laser stimulation could induce CAP in congenitally hearing-impaired mice with a synaptic transmission deficiency between inner hair cells and spiral ganglion neurons. Xia et al. [34] reported a higher neural response when the laser beam targeted spiral ganglion neurons compared to the cochlear wall, while cochlear pressure remained almost independent of the laser irradiation site [34, 36]. These findings suggest that laser stimulation bypasses the need for hair cell depolarization, directly eliciting a cochlear nerve response. This aligns with our earlier results [14, 15], indicating cochlear microphonics and CAP recorded with clicking sounds, whereas pulsed laser irradiation induced only CAP, as reported previously [21, 32]. Additionally, our current results on stimulus generalization show that laser stimulation produces a partially different perception from clicking sounds (Fig. 3I), although the tonotopic response in the inferior colliculus with laser stimulation might correspond to that with clicking sounds when laser irradiation otoacoustically induced the cochlear response [35]. Future studies are essential to investigate the physiological mechanism of laser-evoked auditory perception by comparing the effects of laser irradiation at different levels of peripheral hearing impairment.

### The clinical utility of transtympanic laser stimulation

The clinical utility of laser stimulation depends on whether the laser directly activates auditory nerves or indirectly stimulates them through hair cell activation. If the laser can directly activate auditory nerves, these techniques could pave the way for a groundbreaking hybrid auditory prosthesis, merging laser-evoked perception with residual hearing. Traditional hybrid cochlear implants aim to deliver high-frequency electric hearing while preserving low-frequency residual acoustic hearing [3, 4]. However, hearing preservation has become an important topic of the hybrid cochlear implants due to the complications of post-surgical implantation [3, 5]. Noncontact laser stimulation potentially mitigates this risk. In this study, we investigated the interactions between laser-evoked and auditory-evoked responses by manipulating the relative timing of each stimulation. The data revealed a robust evoked response when both stimuli were presented simultaneously, suggesting that the neural response from transtympanic laser stimulation can be integrated with that from auditory stimulation (Fig. S6). Additionally, simultaneous presentation of auditory and laser stimulation with sub- or near-threshold intensity induced nonlinear facilitation of neural and behavioral responses (Fig. 4). These findings indicate that concurrently presenting auditory and transtympanic laser stimulation has the potential to enhance hearing sensation, offering a potential alternative to traditional hybrid cochlear implants.

**Fig. 4.**
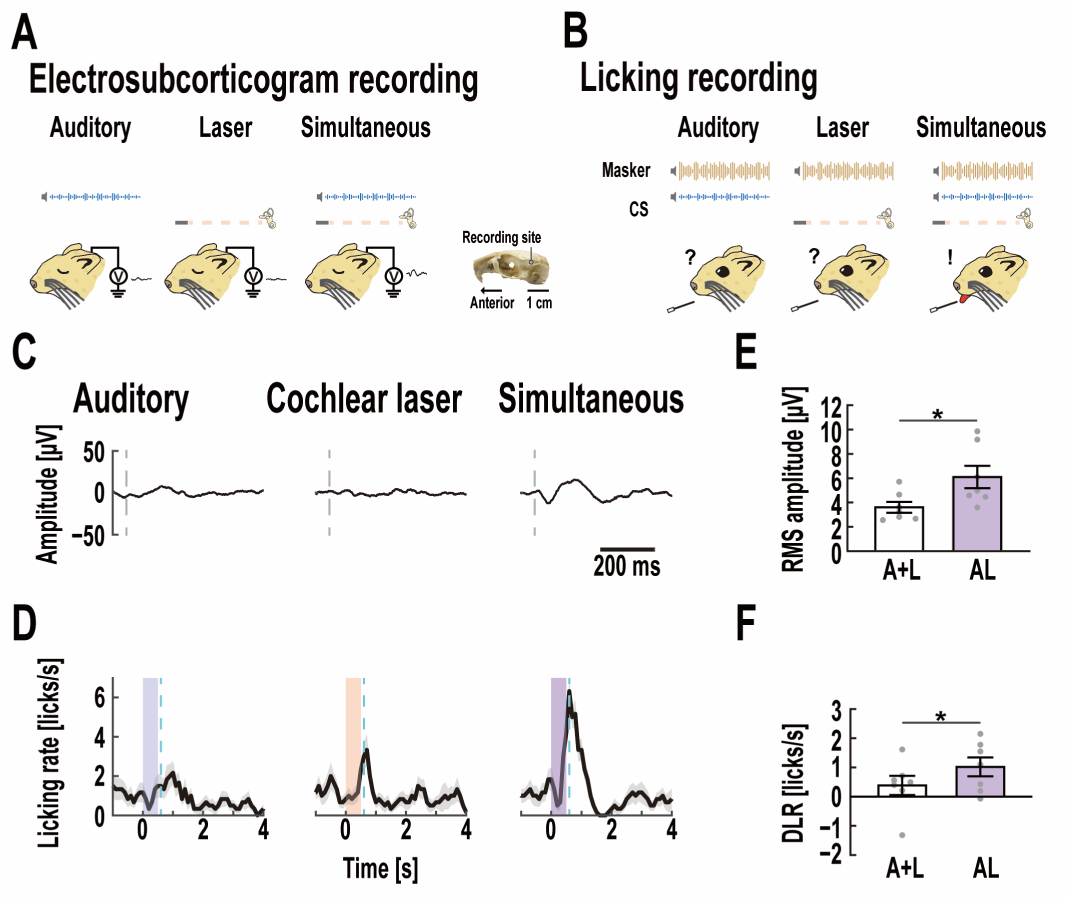
Facilitation of auditory evoked potential (AEP) and behavioral response by simultaneous presentation of auditory and laser stimulation with sub- or near-threshold intensity. Experimental workflow of the Electrosubcorticogram (A) and licking (B) recording are described (Materials and Methods for details). Comparison of AEP (C) and PSTH of licking behavior (D) elicited by auditory, laser, and simultaneous presentation of auditory and laser stimulation. In (C), the gray dotted lines indicate stimulus onset. The blue dotted lines in (D) show the expected reward timing. Gray areas show the SEMs of each trial. The blue, red, and purple zones indicate the auditory, laser, and simultaneous stimulation periods, respectively. Comparing the response elicited by auditory and laser stimulation alone with that by simultaneous stimulation revealed that the latter elicited greater auditory cortical and behavioral responses. Response amplitude of linear summation of auditory- and laser-evoked responses and simultaneous stimulation in AEPs (E) and mean PSTHs of licking behaviors (F). The amplitude evoked by simultaneous presentation in both neural and behavioral response was significantly higher than the amplitude of linearly summating response evoked by auditory and laser stimulation alone (neural response: linear summation (A+L) against simultaneous stimulation (AL), 6.39±1.36 µV *vs.* 8.76±1.23 µV, \**P*<0.05, n=8; behavioral response: linear summation against simultaneous stimulation, 0.39±0.33 licks/s *vs.* 1.02±0.32 licks/s, \**P*<0.05, n=8; Mean±SEM; two-tailed paired *t*-test); the simultaneous stimulation induced a nonlinear facilitation of responses. AEPs and mean PSTHs of licking behavior elicited by auditory and laser stimulation with suprathreshold intensity are shown in Fig. S7.

Even if the laser-evoked CAP is mediated by hair cells rather than direct stimulation of the cochlear nerve, transtympanic laser stimulation could still be advantageous as a middle ear implant. Middle ear implants are commonly employed for patients with moderate to severe hearing deficiencies that cannot be adequately addressed with hearing aids [44]. While middle ear implants can convey greater hair cell vibrations to the inner ear [45, 46], they necessitate surgical implantation with potential complications akin to cochlear implants [47]. Our data demonstrate that the simultaneous presentation of auditory and laser stimulation produces a more pronounced neural and behavioral response than either stimulation alone (Fig. 4 and Fig. S7). Considering that transtympanic laser stimulation bypasses middle ear function [14], it can enhance hair cell activity, potentially amplifying the sensation of hearing. Although further experiments are needed, especially to elucidate the physiological mechanisms of cochlear laser stimulation, our data indicate the feasibility of a laser auditory prosthesis. Transtympanic laser stimulation alone could serve as a viable alternative to electric stimulation in cochlear implants and may prove useful in amplifying hearing sensation by simultaneous presentation of auditory and laser stimulation.

## Conclusion

In summary, this study explored the potential of infrared laser stimulation to reduce the invasiveness of surgery associated with cochlear implants by leveraging its contactless feature. While many studies have concentrated on intracochlear stimulation system to enhance the spectral resolution of cochlear implants using laser stimulation [9–11] or optogenetic stimulation [12, 13], none have investigated the potential of infrared laser stimulation for minimizing surgical invasiveness, considering its contactless nature [14, 15]. Our data provides the first demonstration that laser irradiation of the cochlea from the outer ear can elicit a distinct behavioral auditory response. Laser-conditioned animals successfully learned licking behavior, with conditioned responses and behavioral properties comparable to auditory-conditioned animals. The laser-evoked response was significantly inhibited by auditory masking, and auditory-conditioned animals demonstrated stimulus generalization to laser stimulation. In a subsequent experiment, simultaneous presentation of auditory and laser stimulation induced nonlinear amplification in both auditory cortical and behavioral responses, suggesting that combining auditory and laser stimulation can enhance auditory perception beyond the effects of each stimulus alone. These findings indicate that infrared laser irradiation of the cochlea can potentially evoke and enhance auditory perception, making it a promising candidate for implementation in auditory prostheses.

## Acknowledgements

We would like to thank Hiroshi Riquimaroux for his support during the early stages of this work. We also thank Steffen Hage for technical support and valuable discussion and Yee Ping Cheung and Enago (https://www.enago.com/) for English language editing.

## Author contributions

Conceptualization: Y.T. and K.I.K.; Data curation: Y.T., M.U., A.O., and K.H.; Formal analysis: Y.T. and M.U.; Funding acquisition: Y.T., K.T., and K.I.K.; Investigation: Y.T. and M.U.; Methodology: Y.I., K.T., and S.H.; Project administration: Y.T., K.T., S.H., and K.I.K.; Software: Y.T. and M.U.; Supervision: Y.T., K.T., S.H., and K.I.K.; Validation: Y.T., M.U, and A.O.; Visualization: Y.T.; Writing-Original Draft: Y.T. and M.U.; Writing-Review&Editing: Y.T., K.T. S.H., and K.I.K.

## Data and materials availability

All data needed to evaluate the conclusions are shown in the paper and supplemental figures. Additional data related to this paper are available from the corresponding author upon reasonable request.

## Supplementary Materials

## Materials and methods

### Animals and their maintenance

In total, fifty-seven experimentally naive adult Mongolian gerbils (twenty-six females, thirty-one males; *Meriones unguiculatus*) aged 2–9 months were utilized. Each gerbil was housed with two to four others in a cage measuring 20 cm (W) × 40 cm (L) × 17 cm (H), with free access to food and water. The animal room maintained a temperature between 22°C and 23°C, with approximately 50% relative humidity and a 12-h light: 12-h dark cycle. All experimental procedures adhered to the guidelines established by the Ethics Review Committee of Doshisha University. The Animal Experimental Committee of Doshisha University approved the experimental protocols (reference number: A22071).

### Animal surgery

All surgical procedures were conducted under anesthesia using an intramuscular combination injection of ketamine (47.0 mg/kg) and xylazine (9.3 mg/kg). Breathing pattern was monitored throughout the surgery to assess the depth of the anesthesia. Maintenance doses of ketamine (17.5 mg/kg) and xylazine (7.0 mg/kg) were administered every 30–50 minutes or if the animal exhibited faster respiration rates. Gerbil body temperature was maintained using heating pads positioned beneath the animals.

The parietal and temporal sides of the subject’s head were exposed after shaving the fur. Both sides of the temporal muscle were excised following the application of a local anesthetic (xylocainegel; Aspen Japan, Tokyo, Japan). The skull was meticulously cleaned with a 0.3% sodium hypochlorite solution and saline. A small hole (diameter: 1 mm) was drilled into the temporal skull, located 3.0 mm posterior from the bregma and 4 mm ventral to the lambda–bregma plane, for a recording electrode. Another small hole was made in the dorsal skull, 2 mm anterior to the bregma and 1 mm lateral to the midline, for reference. Reference and recording electrodes (Nilaco, Tokyo, Japan; diameter: 0.13 mm; impedance <20 kΩ) were inserted into these holes and securely fixed at a depth of approximately 1 mm from the cortex surface with acrylic glue. Electrode insertion sites were determined based on the brain atlas of Mongolian gerbils [1] and our prior study [2, 3]. Following the temporal fixation of electrodes with acrylic glue, the entire exposed skull was covered with dental resin (Super bond C&B; Sun Medical, Shiga, Japan). A V-shaped metal plate (F-911; Hilogik, Osaka, Japan) was attached to the dental resin over the skull using dental cement (Province; Shofu, Kyoto, Japan), allowing the subjects to be head-fixed during the experiment.

The left side of the pinna was detached to provide a clear view of the tympanic membrane and outer canal. An incision of the muscle and skin from the shoulder to the jaw exposed the left side of the bulla, and a 1-mm diameter hole was made on the bulla. A silver electrode for recording cochlear responses (Nilaco, Tokyo, Japan; diameter: 0.13 mm; impedance <20 kΩ) was inserted into the hole and secured onto the bony rim of the round window. The electrode hole was completely sealed using acrylic glue and dental cement (Provinice; Shofu, Kyoto, Japan) to stabilize the electrode with the bulla and preserve moisture in the middle ear. All incised muscles and skin were carefully sutured after providing a mixture ointment of colistin sulfate and bacitracin (dolmaisin ointment; Zeria, Tokyo, Japan) as postoperative pain management.

After surgery, all subjects were individually housed and given a recovery period of at least two days before training commenced. The animal surgery followed the same procedure as in our previous studies [4, 5].

### Auditory stimuli

Click trains with a repetition rate of 4 kHz served as auditory stimuli. The click train comprised 0.1 ms rectangular pulses, and the duration of auditory stimulation was 500 ms. The auditory stimulus was delivered through a speaker (AS01008MR-2-R, Pui audio, Ohio, America), positioned 4.5 cm vertically downward at a 45° angle from the gerbil’s left ear. Auditory stimulus generation employed a digital-to-analog converter (Octa-Capture, Roland, Shizuoka, Japan) with a sampling rate of 192 kHz. Signal amplification was achieved using an amplifier (A-10, Pioneer, Tokyo, Japan). The sound pressure level was adjusted within the range of 20–95 dB SPL using a microphone (Type 7016, Aco, Tokyo, Japan).

### Laser stimuli

A repetitive pulsed infrared laser was employed for the infrared laser stimuli. Each individual laser pulse had a rectangular shape in intensity, lasting 0.1 ms, and the inter-onset interval was 0.25 ms, corresponding to a 4-kHz repetition rate. The duration of the pulsed infrared laser train was 500 ms. The pulsed infrared laser was generated using a continuous mode diode laser stimulation system (BWF-OEM, B&W TEK, Delaware, USA) with a wavelength (λ=1871) consistent with previous studies [3–8]. Voltage commands for the laser stimuli were generated using a digital-to-analog converter (Octa-Capture, Roland, Shizuoka, Japan). The laser was delivered through an optical fiber (diameter: 100 μm; NA: 0.22) inserted mediolaterally into the left outer ear canal, with the tip angled rostrally by 10° and dorsally by 5° using a micromanipulator (MM-3; Narishige, Tokyo, Japan) in awake head-fixed subjects. The optic fiber tip was fixed at approximately 0.7 mm (the distance between the cochlea and fiber tip was about 2.2 mm) in front of the tympanic membrane to ensure transtympanic irradiation of the cochlea from the outer ear, as described in our previous reports [3–5, 8]. The laser beam profile at the cochlea (2.2 mm from the fiber tip) was measured, as in previous studies [4, 5]. Radiant energy was measured using a digital power meter (PM100D; Thorlabs, Tokyo, Japan) with a thermal power sensor (S302C; Thorlabs, Tokyo, Japan), calibrated between 0.1 and 13.2 mJ/cm^2^. The beam width (full width at half the maximum diameter) at the cochlea was calculated using an InGaAs-biased detector (DET10D/M; Thorlabs, Tokyo, Japan) and measured as 2.2 mm.

### Visual stimuli

A white light-emitting diode (LED) (diameter: 5.0 mm) was used for visual stimulation. The LED was placed 10 mm from the left eye of the gerbil in the horizontal plane. The voltage command for the visual stimulus was controlled using a digital-to-analog converter (Octa-Capture, Roland, Shizuoka, Japan) with a 192-kHz sampling rate. Light stimulation was applied for 500 ms, and light intensity was calibrated at 96 lux using a digital lux meter (GL-03; be-s, Osaka, Japan).

### Synchronous presentation of auditory and laser stimulation

Simultaneous presentation of click train and pulsed laser train was performed to assess the feasibility of simultaneous presentation of auditory and laser stimulation. The stimulus features of simultaneous stimulation were the same as those of auditory and laser stimulation.

### Apparatus used in the behavioral experiment

Behavioral experiments and electrophysiological recording were performed in the same 60 × 60 × 60-cm (W × L × H) soundproof box, following our previous study [5]. Each subject was positioned on a custom-designed covered platform with a copper sheet. The subjects’ heads were stabilized by clamping the V-shaped metal plate attached to the heads. A metal drinking spout was fixed around 1 mm in front of the mouth of the subjects. A touch sensor (DCTS-10; Sankei Kizai, Tokyo, Japan), connected between the drinking spout and copper sheet of the stage, recorded the timing of licks at 2000 samples/s. Voltage changes from the touch sensor (licking timing) were captured by Arduino Uno (Arduino, Ivrea, Italy) using Python 3.7 on Spyder 501 and stored on a computer. All behaviors were recorded using a video camera (HD-5000; Microsoft, Washington, USA).

### Procedure of the training and test session

After at least two days of recovery from surgery, the gerbils underwent water restriction in their home cage. The training session typically consisted of 250–400 trials per day. The head-fixed animals could voluntarily lick the spout throughout the behavioral experiment. A 3.2 µl drop of water (unconditioned stimulus: US) was delivered through the tube after a short interval, 678±32 ms (mean±SD, n=120), from the onset of auditory, laser, or visual stimulation (i.e., conditioned stimulus: CS). The inter-trial interval was randomly varied between 7 and 13 s. The timing of US and CS was controlled using MATLAB (MathWorks, Massachusetts, USA) and a microcontroller (Arduino Uno; Arduino, Ivrea, Italy) with a custom-made relay circuit.

After animals exhibited elevated licking rates following the CS without the delivery of the US, a test session was conducted. The test session consisted of 280 rewarded trials and 120 nonrewarded trials (i.e., catch trials). In rewarded trials, the CS and US were delivered as in the training session, aiming to sustain a heightened motivational state in the subjects. Catch trials evaluated the subjects’ perceptions without reward feedback. Six stimulus types were presented 20 times in catch trials, and the behavioral response to each stimulus was examined. Additional details for each stimulus condition are provided in Figs. S1 and S2. Changes in licking behavior with conditioning are shown in Fig. S3.

### Recording and analysis of behavioral data

An averaged peri-stimulus time histogram (PSTH) of licking behavior assessed the time course of behavioral responses. PSTHs were calculated in a 100-ms bin size. To smooth the PSTHs, a three-point moving average (i.e., 300 ms window) was applied. To quantify behavioral response, peak licking rates, durations, peak latency, response latencies, and delta licking rate (DLR) were extracted from PSTHs, as in our previous study [5]. Peak licking rates and latency were defined as the licking rate and the time when the licking rate reached its maximum value within 4 s after stimulus onset, respectively. The baseline licking rate was determined as the prestimulus licking rate obtained for 1 s, and duration was quantified as the temporal length of PSTH >50% of the licking rate from the mean baseline licking rate to the peak licking rate. Response latency was calculated by measuring the time between stimulus onset and the behavioral response reaching the mean plus three standard deviations of the baseline licking rate. The DLR was defined as the difference between the mean baseline licking rate and the post-stimulus licking rate (analysis range: 0.2–1.2 s) to evaluate the response amplitude. The analysis range of post-stimulus licking behavior was determined by latency data of the behavioral response (Fig. S5) and our previous study [5].

### Electrophysiological recording during the behavioral experiment

The cochlear response during the behavioral experiment was recorded and amplified 1000 times using a bio-amplifier (MEG-1200; Nihon Kohden, Tokyo, Japan). Electrophysiological data were stored on a computer using an analog-to-digital converter (Octa-capture; Roland, Shizuoka, Japan) at a sampling rate of 96 kHz.

The recorded signal underwent processing with a band-pass filter, employing 1024 digital sampling points (500–3500 Hz) to minimize summating potentials and cochlear microphonics (CMs) while extracting the compound action potential (CAP) of the cochlear nerves. CAP amplitude was defined as the peak-to-peak voltage between the first minimum (N1) and maximum (P1) of each CAP response.

In the auditory masking experiment, defining the peak of CAP became challenging due to the substantial CM elicited by loud white noise (i.e., masker) compared to CAP. The band-pass filter could not entirely eliminate CM. Consequently, the change in the root mean square (RMS) amplitude between pre- and post-signal onset was measured (i.e., diff RMS amplitude). Signal onset was determined through visual inspection using MATLAB (MathWorks, Massachusetts, USA). Pre- and post-CAP signals for 2 ms from signal onset were obtained to calculate the relative voltage change in CAP triggered by auditory and laser stimulation.

### Auditory-evoked potential (AEP) recorded from anesthetized subjects

To explore the impact of synchronous auditory and laser stimulation on the central nervous response, AEPs were recorded from the right auditory cortex using an inserted silver electrode (Nilaco, Tokyo, Japan; diameter: 0.13 mm; impedance <20 kΩ) in anesthetized animals with ketamine (47.0 mg/kg; i.m.) and xylazine (9.3 mg/kg; i.m.). The AEP signal was amplified by a factor of 1000 using a bio-amplifier (MEG-1200; Nihon Kohden, Tokyo, Japan) and stored on a computer through an analog-to-digital converter (Octa-capture; Roland, Shizuoka, Japan) at a sampling rate of 192 kHz. Maintenance doses of ketamine (17.5 mg/kg; i.m.) and xylazine (7.0 mg/kg; i.m.) were administered every 30 min during recording. Subjects were placed on the same covered elevated platform in the soundproof box as in the behavioral experiment, with portable warm packs attached to the cover and small pads beneath to maintain temperature.

To investigate laser interactions with the auditory-evoked response, both clicking sound (35 dB pe. SPL) and pulsed laser (1.1 mJ/cm^2^) were presented, varying the relative timing by 0, 5, 10, or 20 ms before or after laser stimulation. Clicking sound and pulsed laser were generated using a digital-to-analog converter (Octa-Capture, Roland, Shizuoka, Japan) with a 192 kHz sampling rate. AEP signals between 50 and 300 ms from the laser stimulus onset were captured, and the RMS of the extracted signal was calculated as the response amplitude of the AEP signals. The analysis range, determined based on previous studies [9–13], aimed to evaluate auditory cortical activity correlating with auditory perception and avoid the analysis of auditory stimulus artifacts. The change in AEP based on the relative timing of stimulus onset is illustrated in Fig. S6.

Furthermore, a simultaneous presentation of click trains (20, 30, 40, and 50 dB SPL) and pulsed laser trains (0.1, 0.2, 1.1, and 6.9 mJ/cm^2^), each lasting 500 ms with a repetition rate of 4 kHz, was conducted to evaluate the feasibility of concurrent auditory and laser stimulation. These sound pressure levels and radiant energy values were chosen to cover sub- to suprathreshold intensities [2–4, 8, 14]. AEP responses elicited by the simultaneous presentation of auditory and laser stimulation, resulting in nearly equal AEP amplitude (i.e., higher AEP amplitude induced by auditory or laser stimulation not exceeding twice the lower response amplitude), were recorded. The extent of nonlinear amplification based on the response amplitude induced by the mono stimulus was investigated. The response amplitude induced by mono-stimulation was defined as the median of the response amplitude induced by auditory and laser stimulation alone. Data were categorized based on the response amplitude induced by mono-stimulation, as the nonlinear amplification tendency varied notably between groups where no observable response (amplitude: 3.5 µV or lower) and conspicuous response (amplitude: 20 µV or higher) were observed with mono-stimulus presentation. Mono-stimulus intensities that elicited no observable response and conspicuous responses were defined as sub- or near-threshold and suprathreshold intensities, respectively.

## Supplemental Figure

**Fig. S1.**
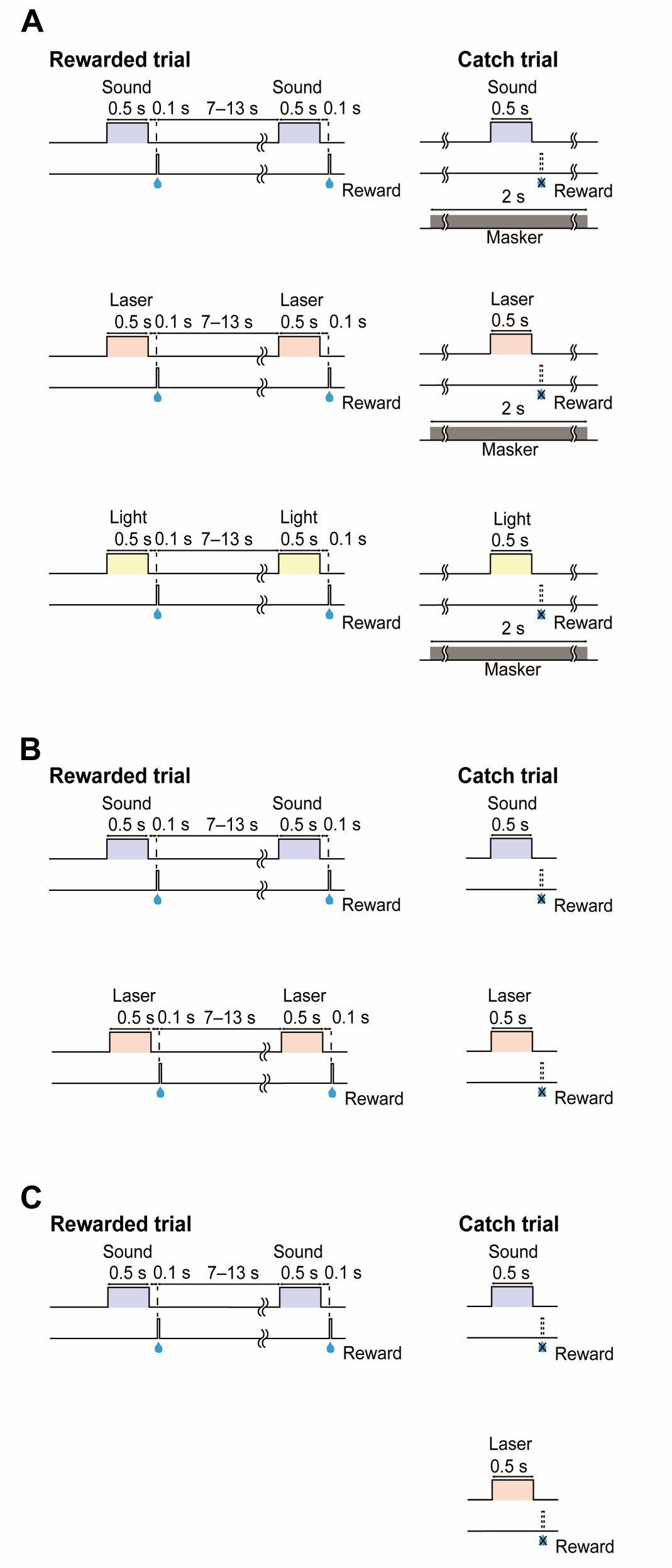
Schematic of conditioned (CS) and unconditioned (US) stimulus presentation in the mono-stimulus experiment. The behavioral task of effect of auditory masking on auditory- and laser-evoked responses (A), intensity dependance of auditory- and laser-evoked responses (B), and assessment of stimulus generalization (C) are shown. The timing of CS and US in the rewarded (left) and catch trials (right) are described. In (A), 15 auditory (70-dB SPL)- and 6 laser (11.7 mJ/cm^2^)-trained animals were used. In the rewarded trial (left), click train (70 dB SPL) or pulsed laser train (11.7 mJ/cm^2^) was presented to the auditory- and laser-trained animals, respectively. In the catch trials (right), the sound pressure level and radiant energy of CS were varied (auditory: 20, 35, 50, 65, 80, or 95 dB SPL; laser: 0.1, 1.1, 3.1, 6.9, 10.2, or 13.2 mJ/cm^2^). In (B), the experiment used six auditory-(80 dB SPL), six laser-(13.2 mJ/cm^2^), and four visual-(96 lux) trained animals. In the rewarded trial, click train (80 dB SPL), pulsed laser train (13.2 mJ/cm^2^), or LED light (96 lux) was presented to the auditory-, laser-, or visual-trained animals, respectively. To investigate the effect of auditory masking on the physiological and behavioral responses induced by auditory, laser, and visual stimulation, white noise (700–44,800 Hz) of 0, 35, 50, 65, 80, or 95 dB SPL for 2 s was presented during the auditory, laser, or visual stimulus periods of catch trials. In (C), all animals (n=11) were trained by auditory (70-dB SPL) stimulation. In the rewarded trials, a click train was used. Auditory (20, 50, and 80-dB SPL) and laser (0.1, 6.6, and 13.2 mJ/cm^2^) stimuli were introduced in the catch trials.

**Fig. S2.**
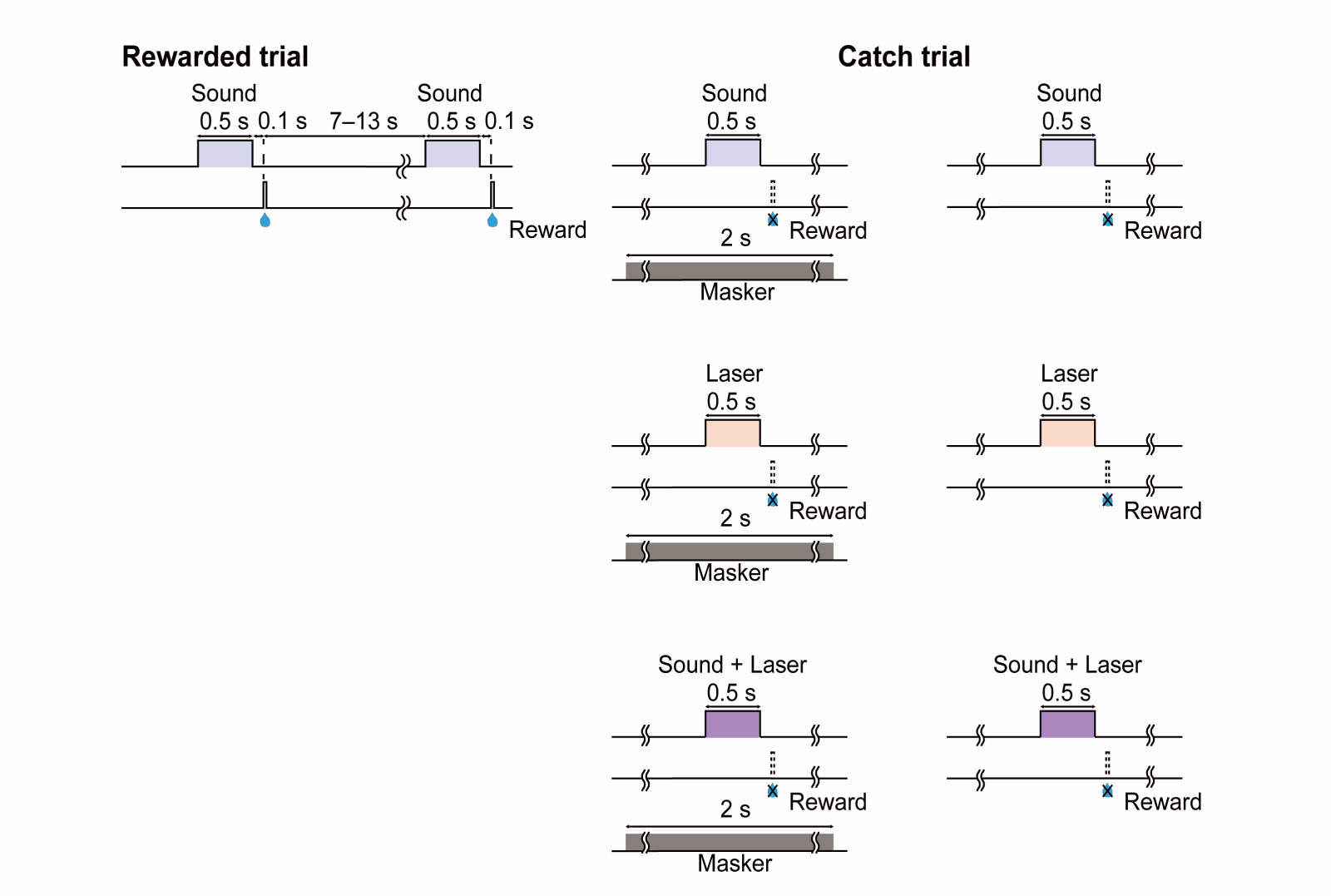
Illustration of behavioral tasks for evaluating the feasibility of simultaneous presentation of auditory and laser stimulation. The timing of conditioned and unconditioned stimuli in the rewarded (left) and catch trials (right) are described. Eight subjects were trained with auditory (80-dB SPL) and laser (13.2 mJ/cm^2^) stimulation. White noise (70 dB SPL) was presented during the stimulus period in the same way as in the auditory masking experiment to assess the efficiency of simultaneous stimulation in a quiet and noisy environment. An auditory (80-dB SPL) stimulus was introduced in the rewarded trials. In the catch trials, auditory (80 dB SPL), laser (13.2 mJ/cm^2^), or simultaneous stimulation was presented with or without background white noise.

**Fig. S3.**
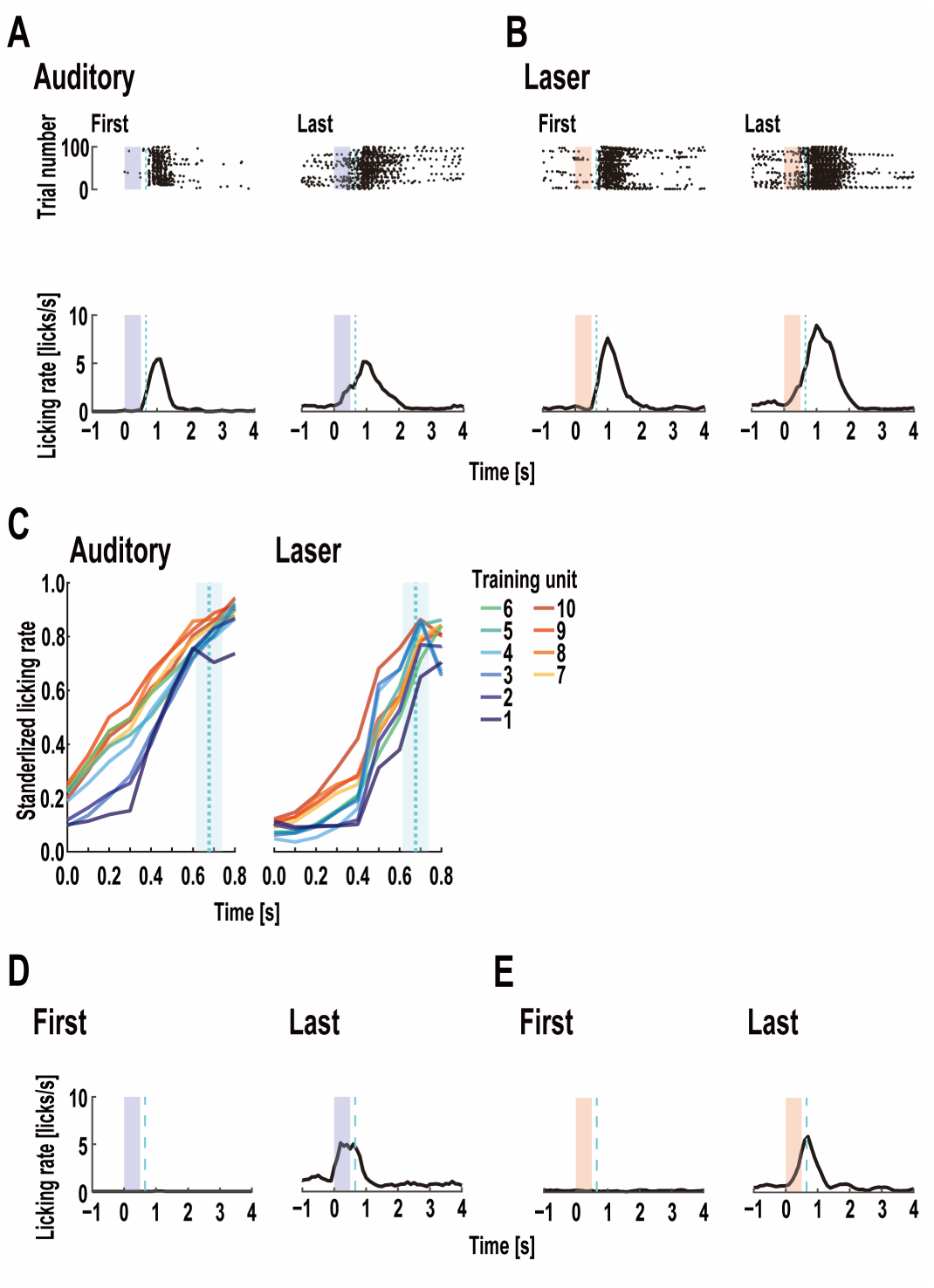
Licking behavior transition by Pavlovian conditioning with auditory and laser stimulation. Auditory-(A) and laser-induced (B) licking behavior in the first (1– 100 trials) and last (901–1000 trials) training units. Individual trials (top) and mean peri-stimulus time histogram (PSTH) of licking behavior (bottom) elicited by auditory and laser stimulation are described. The blue and red zones indicate the auditory and laser stimulation periods, respectively. The blue dashed lines indicate the reward timing. The black plots in the top figure show the licking timing. In the first training unit, auditory- and laser-evoked licking responses were observed after US presentation, while these licking behaviors were also recorded before US presentation in the last training unit. (C) Changes in average auditory (n=13)- and laser (n=12)-CR in each training unit. One training unit comprised 100 trials. The blue dashed lines and areas show the mean reward timing and 2SD. The standardized licking rates until 0.6 s after stimulus onset were measured as CR. The gradual increase in CR amplitude was observed every session. Auditory-(D) and laser-induced (E) average PSTH of licking rates on the first and last day of catch trials (1–120 trials). The blue and red zones indicate the auditory and laser stimulation periods, respectively. The blue dashed lines indicate the reward timing in the rewarded trial. Catch trials were performed to monitor the association between CS and US. Synchronous increase in licking rates with auditory and laser stimulation were not obtained on the first day of the catch trial. After completing the training, synchronous licking behaviors with CS were observed.

**Fig. S4.**
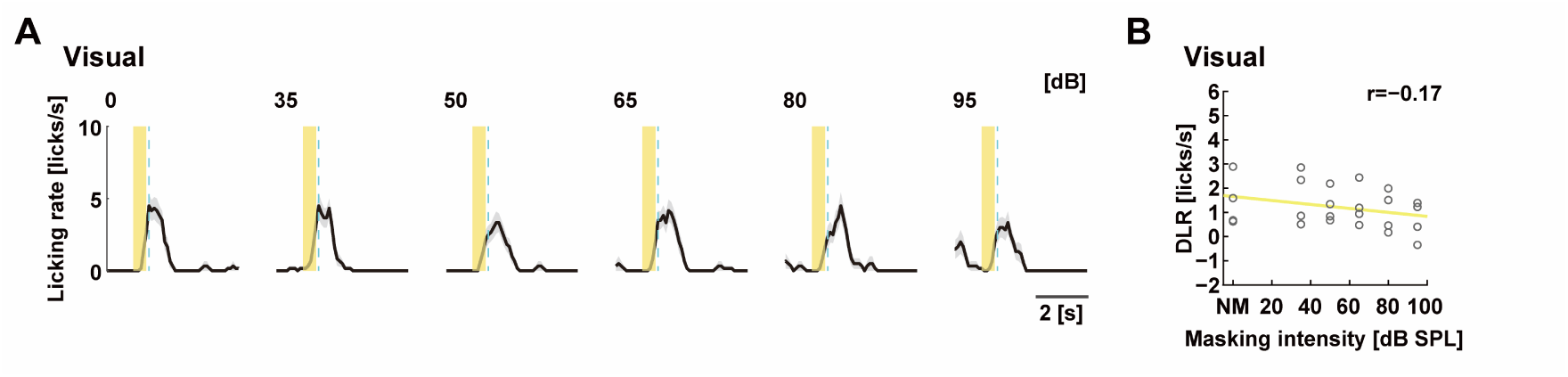
Effect of auditory masking on visually conditioned response. This investigation was performed as a control condition to ensure that the presence of intensive white noise did not cause behavioral disturbance or interfere with auditory input. (A) Mean PSTH of licking behavior elicited by visual stimulation in the masking experiment. Yellow areas indicate stimulus duration. Blue dashed lines indicate reward timing in a training session. (B) Correlation between masking intensity and visually evoked DLR. The result of the control experiment showed that a significant decrease in visual-evoked DLR was not observed as the sound pressure level of white noise increased (r=-0.17, p=0.310; Pearson’s correlation analysis); therefore, auditory masking in this experiment mainly interrupted auditory-related perception.

**Fig. S5.**
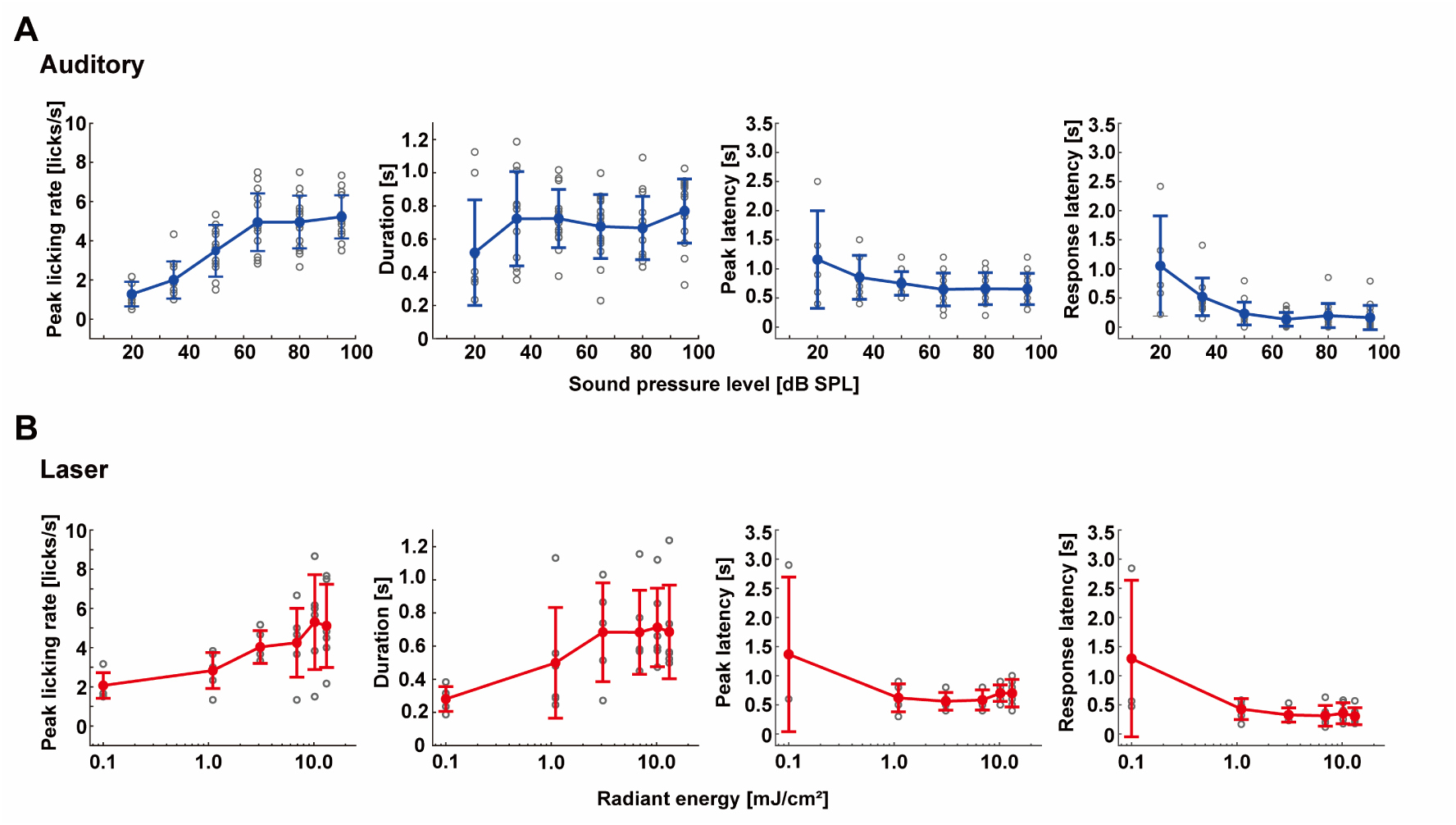
Changes in features of behavioral responses by modulating sound pressure level and radiant energy. Intensity dependance of the peak licking rate (leftmost), duration (left), peak latency (right), and response latency (rightmost) with auditory (A) and laser (B) stimulation. Error bars show the standard deviations. As the sound pressure level increased from 20 to 95 dB SPL, the peak licking rate rose from 1.28±0.63 to 5.22±1.10 licks/s; duration increased from 0.52±0.32 to 0.77±0.19 s; peak latency decreased from 1.16±0.84 to 0.65±0.27 s; and response latency decreased from 1.05±0.86 to 0.16±0.21 s. As radiant energy increased from 0.1 to 13.2 mJ/cm^2^, the peak licking rate rose from 2.07±0.65 to 5.11±2.13 licks/s; duration increased from 0.28±0.07 to 0.69±0.28 s; peak latency decreased from 1.37±1.33 to 0.70±0.24 s; and response latency decreased from 1.29±1.34 to 0.31±0.15 s. The properties of auditory-evoked behavioral responses were similar to those in previous studies [5] and to those of laser-evoked behavioral responses.

**Fig. S6.**
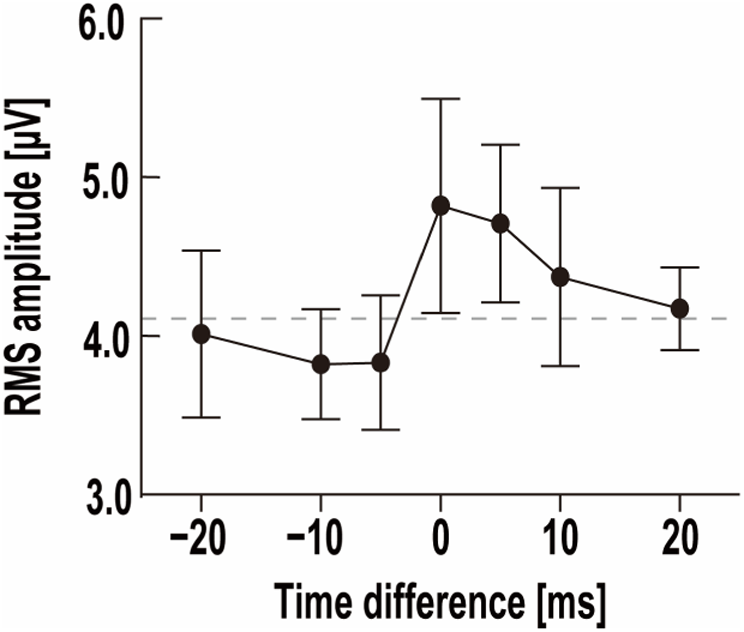
Auditory-evoked potential (AEP) depending on the difference in the relative timing of stimulus onset. The effect of the relative timing of auditory and laser stimulation on AEP and their response amplitudes are shown. The gray dashed line is the baseline amplitude, calculated by averaging the root mean square (RMS) amplitude of the prestimulus electrophysiological signal obtained for 50 ms. When the auditory stimulus was presented before the laser stimulus, the RMS amplitude decreased as the time difference of the stimulus onset increased (from −5 to −20 ms). The peak RMS amplitude was observed at simultaneous stimulation. Then, the amplitude decreased as the auditory stimulation lag increased (5–20 ms).

**Fig. S7.**
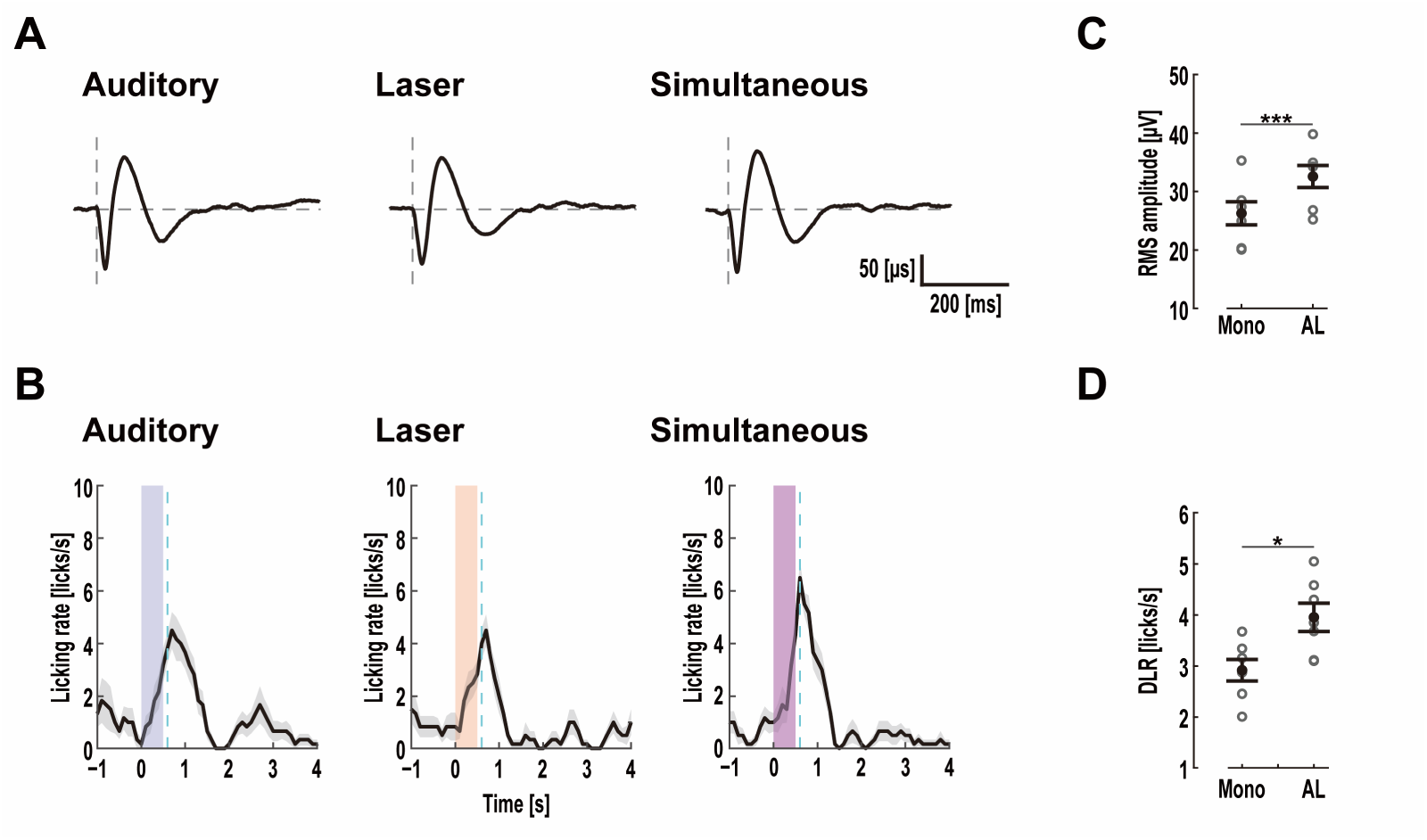
Interaction between auditory and laser stimulation with suprathreshold intensity. Comparison of AEP (A) and PSTH of licking behavior (B) induced by suprathreshold auditory, laser, and simultaneous stimulation. In (A), the vertical and horizontal dashed gray lines show stimulus onset and 0 µV, respectively. The blue dashed lines in (B) show the expected reward timing. Gray areas show the SEMs of each trial. The blue, red, and purple zones indicate the auditory, laser, and simultaneous stimulation periods, respectively. Response amplitude by mono-stimulus and simultaneous stimulation in neural (C) and behavioral (D) responses. When we simultaneously presented suprathreshold auditory and laser stimulation, a nonlinear facilitation was not observed in neural and behavioral responses, unlike simultaneous stimulation with sub- or near-threshold intensity shown in Fig. 4. However, simultaneous presentation of auditory and laser stimulation with supraintensity induced significantly higher response amplitudes than mono stimulation; simultaneous presentation of suprathreshold auditory and laser stimulation induced significantly greater neural and behavioral responses than mono stimulation, but the degree of amplification caused by suprathreshold simultaneous stimulation was less than that caused by sub- or near-threshold simultaneous stimulation.

